# ralphi: a deep reinforcement learning framework for haplotype assembly

**DOI:** 10.1101/2025.02.17.638151

**Authors:** Enzo Battistella, Anant Maheshwari, Barış Ekim, Bonnie Berger, Victoria Popic

## Abstract

Haplotype assembly is the problem of reconstructing the combination of alleles on the maternally and paternally inherited chromosome copies. Individual haplotypes are essential to our understanding of how combinations of different variants impact phenotype. In this work, we focus on read-based haplotype assembly of individual diploid genomes, which reconstructs the two haplotypes directly from read alignments at variant loci. We introduce `ralphi`, a novel deep reinforcement learning framework for haplotype assembly, which integrates the representational power of deep learning with reinforcement learning to accurately partition read fragments into their respective haplotype sets. To set the reward objective for reinforcement learning, our approach uses the classic reduction of the problem to the *maximum fragment cut* formulation on fragment graphs, where nodes correspond to reads and edge weights capture the conflict or agreement of the reads at shared variant sites. We trained `ralphi` on a diverse dataset of fragment graph topologies derived from genomes in the 1000 Genomes Project. We show that `ralphi` consistently achieves lower error rates at comparable or longer haplotype block lengths over the state of the art for short and long ONT reads at varying coverage in standard human genome benchmarks. `ralphi` is available at https://github.com/PopicLab/ralphi.

## 1 Introduction

Haplotype assembly, also known as genome phasing, is the problem of reconstructing the sequence of alleles at sites of genetic variation of each individual chromosome copy in a genome. The specific configuration of variants across homologous genome regions allows us to differentiate between *in cis* variants present on the same chromosome molecule and *in trans* variants present across homologous chromosomes. This information is required to discover compound heterozygous variants, which occur when both homologous copies of a gene contain variants at different positions with a potentially deleterious impact on the function of both gene copies, and play a key role in numerous diseases, such as cancer [47, 44, 20], Charcot-Marie-Tooth neuropathy [42], Ataxia-telangiectasia [22, 46], and deafness [50]. As such, the accurate reconstruction of haplotypes is vital to our interpretation of genetic variation and its role in disease [13, 49]. Haplotype assemblies are also essential to the study of population structure, genetic diversity, mechanisms of inheritance, and genome evolution [23, 28, 3, 11].

Numerous approaches have been developed to date for haplotype reconstruction. Population based statistical approaches [21, 12, 41] infer haplotypes by leveraging observed patterns of genetic variation in a reference panel (e.g. large-scale genotype datasets from the 1000 Genomes Project [19] or the UK Biobank [14]). On the other hand, read-based approaches assemble individual haplotypes using the overlap of read alignments at heterozygous variant loci. Due to sequencing errors, however, the problem of read-based haplotype assembly is computationally intractable. Numerous formulations for this problem have been proposed to date [24, 45, 37, 4, 36, 54, 5, 7, 8, 26, 43, 40]. Among these, some of the most popular definitions include the minimum error correction, minimal fragment removal, and minimal single-nucleotide polymorphism (SNP) removal optimization objectives [48, 45, 24]. Since these formulations are NP-hard and also hard to approximate [17, 9], heuristic-based algorithms are predominantly used by current state-of-the-art methods [26, 43]. However, heuristics cannot tractably capture the full complexity of the confounding factors in this problem domain, such as the diverse error regimes of different sequencing platforms, as well as the non-uniformity of genome coverage and variant density within the genome and across different populations. Learning-based approaches, on the other hand, can detect complex patterns in high-dimensional data, which are too challenging to discover and define manually. Recently, several deep learning methods have been proposed for the problem of haplotype assembly in polypoid species and viral quasispecies [52, 30, 31]. For example, NeurHap [52] formulates haplotype phasing as a graph coloring problem on read-overlap graphs and uses a graph neural network to learn color assignments, while CAECseq [31] uses convolutional auto-encoders to cluster reads into haplotypes. These approaches, however, have been evaluated only on small or partial genomes and read datasets (with most benchmarked datasets containing a few hundred to a few thousand reads and variants); as such, their performance on a full human genome has not been demonstrated.

In this work, we introduce `ralphi`, a novel framework for haplotype assembly based on deep reinforcement learning (DRL), which offers the ability to learn combinatorial optimization algorithms for high-dimensional inputs. To the best of our knowledge, DRL has never been used for haplotype assembly. Briefly, in reinforcement learning (RL), an agent is trained to make decisions in a given environment that maximize its reward through trial and error. DRL integrates the representational ability of deep learning with the trial-and-error-based optimization of RL to enable operations on high-dimensional state spaces. It has been shown to achieve state-of-the-art results in a wide range of combinatorial optimization problems, including Traveling Salesman, Minimum Vertex Cover, and Maximum Cut [32]. In `ralphi`, we train a DRL agent to partition read fragments into two haplotype sets while optimizing the maximum fragment cut (MFC) objective proposed in Duitama et al. [24], which involves solving the NP-hard *weighted max-cut* problem [5, 26, 24] on read-based graphs. `ralphi` leverages a graph convolutional neural network (GCN) to embed fragment graphs and an actor-critic RL model to learn the read-to-haplotype assignment algorithm based on the MFC reward. We used a large set of fragment graph topologies derived from genomes from a diverse set of populations included in the 1000 Genomes Project to train `ralphi`. To demonstrate the generalizability and adaptability of our learning-based approach to multiple sequencing platforms, we generated `ralphi` models for both Oxford Nanopore Technologies (ONT) long-read datasets and Illumina short-read datasets. We demonstrate that `ralphi` predominantly improves the accuracy of the reconstructed haplotypes as compared to the state of the art, while maintaining high phasing block lengths across several standard benchmark datasets from both short and long sequencing platforms and different genome coverage regimes.

## 2 Methods

At a high level, `ralphi` consists of two key modules (Figure 1): (1) fragment graph construction from read alignments and variants, which generates the input to the learning-based framework, and (2) the DRL framework, which partitions graph fragments into two haplotypes by (a) embedding fragment graphs using a GCN, and (b) iteratively assigning each node in the graph to one of the haplotypes using the actor-critic RL algorithm and the MFC reward.

**Fig 1:**
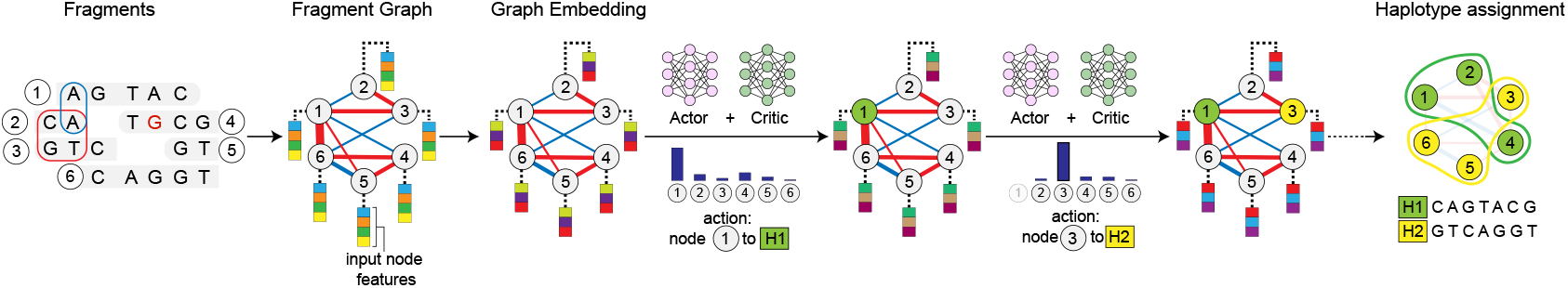
Overview of the Ralphi framework for haplotype assembly. As an example, a fragment graph is constructed from a set of 6 DNA fragments covering 7 SNPs (the allele containing a sequencing error is shown in red), with negative agreement edges shown in blue, and conflict edges, in red (the thickness of the edge corresponds to edge weight). Fragments are partitioned into final haplotypes through an iterative process wherein the agent assigns a node to a haplotype in each step based on the prediction given by the actor-critic network operating over the fragment graph embedding. The final haplotypes are assembled by computing the consensus of the fragments in each partition.

### 2.1 Fragment graph construction

#### Allele matrix

Given a set of read alignments and heterozygous variants, the input to a read-based phasing algorithm is usually represented as a matrix *M* of size *m* x *n*, where *m* is the number of fragments (informative read alignments that span at least two heterozygous variants), *n* is the number of heterozygous variants (e.g. SNPs), and each entry *M*_*i,j*_ stores the allele of fragment *F*_*i*_ covering the locus of variant *j*, with the reference allele usually encoded as 0, the variant allele as 1, and −is used when the fragment does not cover the variant.

#### Fragment graph

As described in [24], a fragment graph *G* = (*V, E*) is constructed from a n allele matrix *M*, such that *V* = {1, …, *m*} and (*u, v*) *∈ E* iff *w*(*u, v*) ≠ 0, where 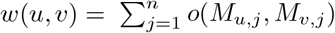 and *o*(*a, b*) is defined as follows. Given two alleles *a* and *b*:

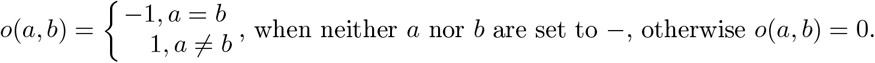

The weight of each edge (*u, v*) in *G* is set to *w*(*u, v*).

As such, each node in a fragment graph corresponds to a fragment *F*_*i*_ in matrix *M* and an edge connects two nodes only if the two corresponding fragments cover at least one common variant. The weight of each edge represents the difference in the number of alleles that the two fragments agree on and disagree on, respectively, such that stronger agreement will result in a negative edge weight, and a stronger conflict will result in a positive edge weight. A cut in this graph naturally represents an assignment of fragments to each haplotype. Intuitively, higher-weight cuts will separate fragments with a higher number of conflicting alleles, which are likely to pertain to different haplotypes.

#### Maximum Fragment Cut (MFC)

The MFC optimization model proposed by Duitama et al. [24] solves the read-based haplotype assembly problem by finding the maximum weighted cut in the fragment graph *G*. Namely, let *C* be a subset of nodes representing a cut of *G*, the fragment cut score is then defined as *s*(*C*) =Σ _*u* ∈ *C*_ Σ_*v* ∉*C*_ *w*(*u, v*). The MFC objective is to find the cut that maximizes *s*(*C*).

Note that fragment graphs are not usually connected, and in practice MFC is computed for each connected component of the graph separately, resulting in an output that typically consists of multiple distinct phased haplotype sets. The accuracy and length of the resulting partial haplotypes in each set are key metrics used to evaluate haplotype assembly as described below.

#### Fragment graph compression

In order to scale to higher coverage datasets, `ralphi` adds an additional graph compression step during construction. In particular, we combine all equivalent fragments into a single node in *G* with an associated compression number *c*, where fragments are considered equivalent if they cover the same set of variants with the same alleles. As a result, edges between equivalent fragments are removed (these are by definition edges that capture perfect fragment agreement and will not be split by an optimal cut) and the weight *w*(*u, v*) of the remaining edges is multiplied by *c*(*u*) \**c*(*v*) corresponding to the compression numbers of the two connected nodes, respectively. By construction, this compression procedure guarantees that the MFC of the compressed and original graphs are equivalent.

#### Pre-processing: read and variant selection

Similar to other read-based phasing tools, `ralphi` performs an initial selection of informative read alignments (provided as an input BAM file) and variants (provided as a VCF file). We primarily rely on previously established best quality control practices for this pre-processing step. For example, `ralphi` performs alignment filtration based on the alignment MAPQ score, as well as read quality for ONT inputs. It also downsamples high-coverage BAM files to minimize runtime using WhatsHap[43], which was shown to outperform random downsampling. To improve the accuracy of alleles observed at variant sites in long-read sequencing, which have a higher occurrence of indel errors, `ralphi` calls into the realignment module of WhatsHap, which realigns a small window of the read around the variant site. To increase fragment contiguity, `ralphi` joins non-overlapping partial alignments of the same read into a single fragment. Finally, given the selected reads, `ralphi` examines the evidence at each variant site and removes variants which are covered only by a single SNP allele, by a single SNP allele and an indel, or only by reads from a single strand (similar to the heuristic used in LongPhase [40]).

### 2.2 Deep reinforcement learning framework for MFC

#### Fragment graph embedding

##### Background

Graph neural networks (GNNs) are a powerful paradigm for graph representation learning and can effectively capture complex combinatorial graph structures [51]. Briefly, GNNs rely on a message-passing scheme, wherein each node aggregates features from its neighbors to update its representation. Starting from an initial set of node features, the representation of each node is updated iteratively using message passing, resulting in a final *z*-dimensional embedding vector. *GNN Architecture:* `ralphi` uses a graph convolutional neural network (GCN) [33] to embed fragment graphs, which consists of a single GCN layer with the ELU activation function [18] and *z* = 264.

#### Input node features

Since the embedding is used to capture the state of the fragment graph after each iteration (i.e., after the assignment of a single node to one of the two haplotypes), each node in the graph is associated with a feature representing its current haplotype assignment. A node can be in three possible states: unassigned, assigned to haplotype 1 (H1), or assigned to haplotype 2 (H2). Additionally, we explored using node features that capture various properties of the graph topology (e.g., node degree), read alignment (e.g., number of covered variants), and algorithmic context (e.g., how many nodes have already been assigned). We found that `ralphi` achieved best results to date when *betweeness centrality* was included as a node feature. Betweeness centrality encodes the fraction of shortest paths passing through a given node *w* and is defined as: 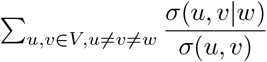, where *σ*(*u, v*) *σ*(*u, v*) denotes the number of shortest paths between *u* and *v*, while *σ*(*u, v w*) is the number of shortest paths from *u* to *v* that pass through *w*. Since computing all shortest paths takes *Θ*(*V* ^3^) time, we use the approximate estimation of betweeness centrality proposed in [10] applied to the unweighted fragment graph. Note, we aim to tune and incorporate additional node features from the categories described above in the next releases of `ralphi`.

#### Reinforcement learning architecture

##### Background

Briefly, in DRL, we define a set of actions *A*, states *S*, and a reward function *R*(*s, a*) that specifies the amount of reward received by the agent when taking an action *a* from state *s*; we then train an agent to learn a policy that maximize its reward through trial and error. *States, actions, and reward:* Based on the max-cut formulation in [32], (1) `ralphi` states correspond to the fragment graph with partial haplotype assignments represented by the graph embeddings, (2) actions involve selecting an unassigned node in the graph and assigning it to one of the two haplotypes, and (3) the reward is given by the change in fragment cut value when assigning a node *u* to haplotype *H*_*k*_, namely: 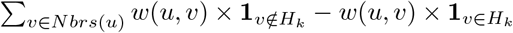, where *Nbrs*(*u*) represents all the neighbors of node *u* in the graph, which have already been assigned to a haplotype. *RL algorithm:* `ralphi` use the popular actor-critic reinforcement learning method [35] to train the agent. This method consists of two networks, the actor and the critic, such that the actor learns which actions the agent should take and the critic learns the value of different actions. In `ralphi`, the actor and the critic are single linear layer networks, with weights initialized using an orthogonal matrix to mitigate the vanishing gradient phenomenon; each layer is also normalized using spectral normalization, which has been shown to increase actor-critic’s stability [16].

#### Training data generation

We generated a diverse set of fragment graph topologies for training, aimed at capturing the variability in genome structure (e.g., variant density within and across genomes in different populations), read depth, sequencing error rates, and read lengths. To that end, we selected the following 10 genomes from the 1000 Genomes Project [15]: HG00234, HG00250, HG00627, HG01598, HG02047, HG03046, HG03166, HG03388, HG04225, and NA11932, which come from diverse populations including British, Southern Han Chinese, Kinh Vietnamese, Gambian Madinka, Esan, Mende, and Telegu. Fragment graphs generated from chromosome 20 were held out for validation. We incorporated the variants of each genome into the GRCh38 reference and generated Illumina short reads and ONT long. For each genome, we generated read datasets at varying depth: 5x, 10x, 15x, and 30x. For short reads, we used the DWGSIM read simulator [1] with 2% and 5% error rates and aligned reads with BWA-MEM [38]. This procedure yielded 3,025,458 graphs, capturing a diversity of populations, coverages, and error rates. For ONT reads, we used nanosim [53] for read simulation and minimap2 [39] for read mapping (using the ONT-specific preset), which yielded 52,360 graphs. Figure 2 highlights the diversity of topologies captured in the resulting dataset. We computed key topological features (e.g., graph size, density, diameter, and number of articulation points) for the resulting graphs and applied graph sampling strategies to balance across different properties, obtaining a training dataset with 79,000 graphs for short reads and 13,091 graphs for long reads. We trained `ralphi` first on short-read graphs, and then fine-tuned the model for ONT reads. We used *curriculum learning* [6] for ONT training, which shows progressively harder examples to the agent as it trains, by ordering our graphs from the smallest and sparsest to the largest and most dense.

**Fig 2:**
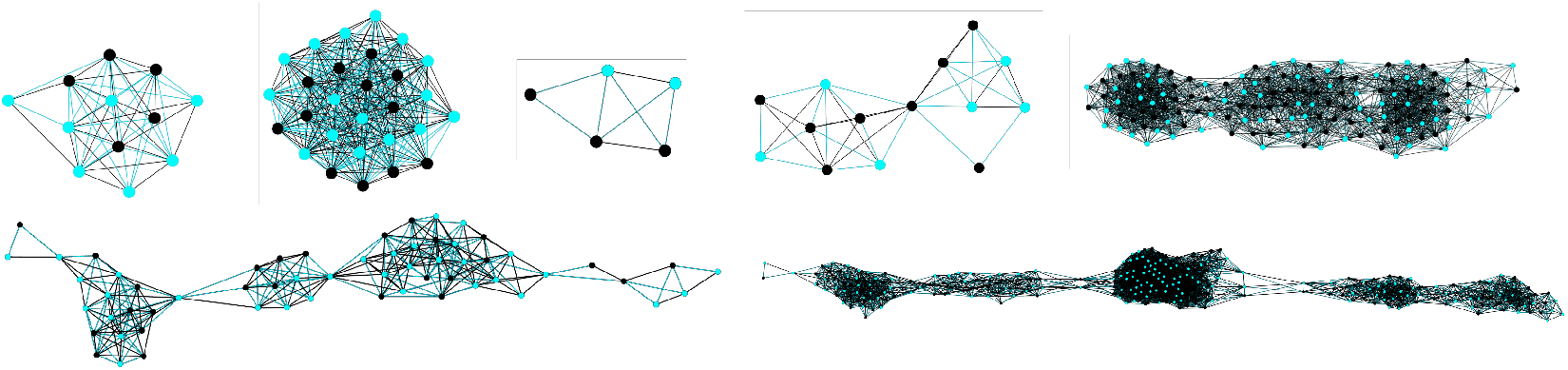
Examples of fragment graph topologies from genomes in the 1000 Genomes Project. Negative and positive edge weights are in black and blue, respectively; node color indicates haplotype membership selected by `ralphi`.

## 3 Results

### 3.1 Evaluation setup

We evaluated the performance of `ralphi` on the NA12878 and HG002 standard benchmark genomes using publicly-available short Illumina reads and long ONT reads. We compared `ralphi` to the popular state-of-the-art phasing methods WhatsHap (2.3) [43], HapCUT2 (1.3.4) [26], and LongPhase (1.5) [40] developed for long reads (note: we used LongPhase only for ONT comparisons; we used version 2.0 of WhatsHap for short reads since it performed better with this older version on this data type). We also included RefHap (1.0.0) to directly compare `ralphi`’s learned algorithm to the heuristic algorithm developed for the same MFC objective (unfortunately only partial metrics are available for RefHap since it did not finish running on all the benchmarks during the course of several months). We ran all the tools in default mode. We could not include a comparison with existing deep-learning methods NeurHap [52] and CAECseq [31], since the available code does not operate with standard phasing inputs and outputs and requires custom model training. To assess the effect of coverage on phasing performance, we downsampled the original read datasets to a depth of 30x, 15x, 10x, and 5x. We compare the following key standard phasing metrics to assess performance: switch error rate, mismatch error rate, and the AN50 score, which assess the accuracy and contiguity of the reconstructed haplotypes, respectively [26]. These metrics were computed using the utility script provided by HapCUT2.

### 3.2 NA12878 benchmark

For the NA12878 benchmark, we used phased high-confidence variants from the Illumina Platinum Genomes callset [25] as ground truth. We obtained Illumina 150-bp paired short reads from the 1000 Genomes Project high-coverage NYGC release [15] and ONT long reads (R10.4.1 chemistry) from the Garvan Institute Long Read Sequencing Benchmark Data platform [27]. Given the variability in read profiles (read length and quality) of ONT sequencing, we also downloaded a second (replica) ONT dataset from the Oxford Nanopore Technologies Open Data project [2] to evaluate performance on the same genome but with different read datasets. Figure 3A shows that `ralphi` consistently achieves lower error rates across all coverage regimes while maintaining or improving on the AN50 haplotype block contiguity metric in both long and short reads. It also achieved the highest AN50 at higher coverage with ONT reads.

**Fig 3:**
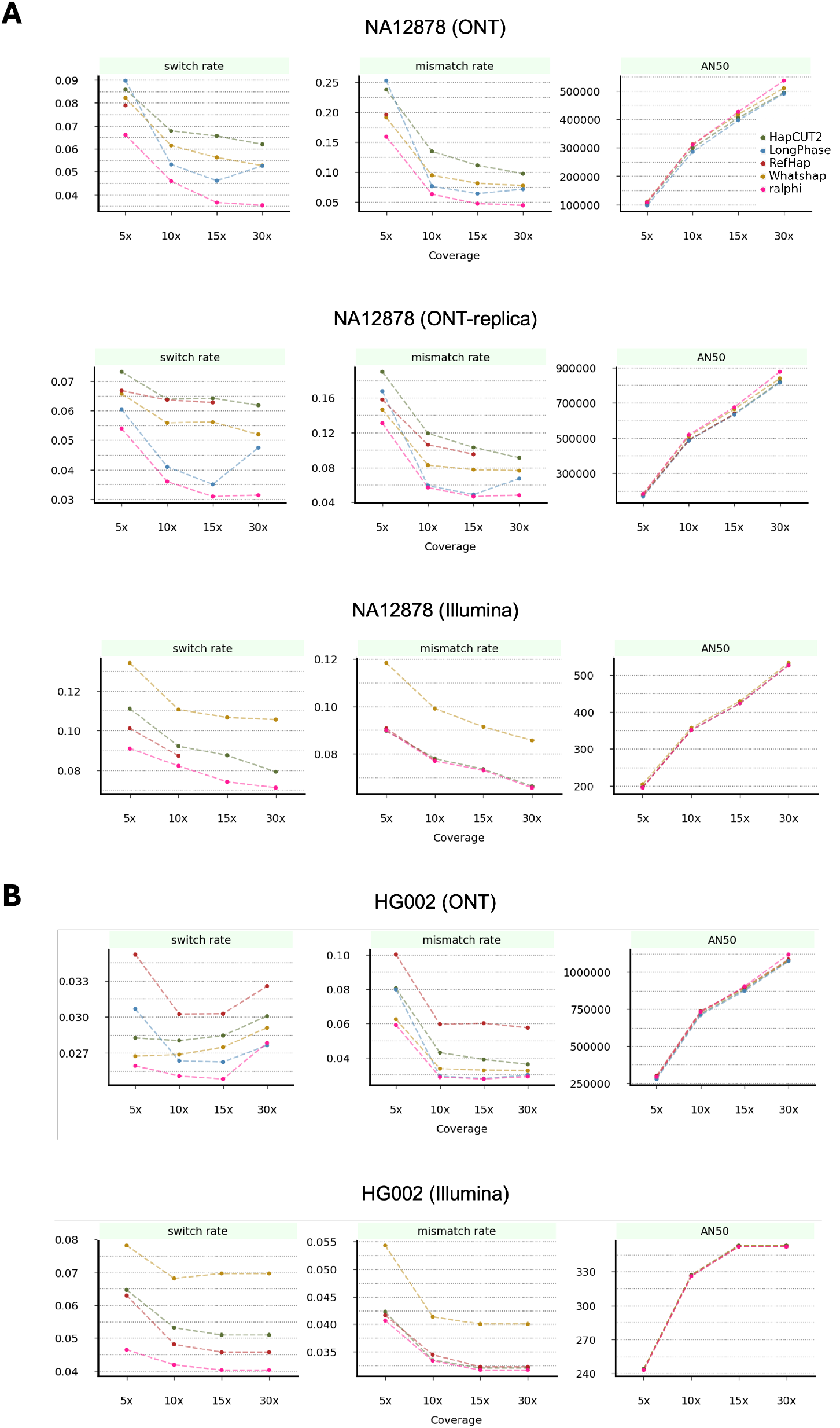
Performance evaluation on the NA12878 (A) and HG002 (B) benchmark genomes using ONT long reads and Illumina short reads. Note: switch and mismatch error rates are a percentage.

### 3.3 HG002 benchmark

For the HG002 benchmark, we used phased variants from the 4.2.1 GIAB release [55] as ground truth. We obtained 250-bp paired Illumina short reads from GIAB [29] and ONT long reads (R10.4.1 chemistry) from the AnVIL workspace [34]. Figure 3B shows that `ralphi` similarly achieved lower error rates across most settings in this benchmark, and a slightly higher AN50 at higher coverage for ONT reads.

#### 3.4 Runtime evaluation

We evaluated the runtime of `ralphi` and other methods on chr1 of the NA12878 genome benchmark. All our experiments were performed on an AMD Ryzen Threadripper PRO 3995WX system with 516GB of RAM. We report the runtime of each tool on a single core for both short reads and long reads across all coverage regimes in Tables 1 and 2. On ONT reads, LongPhase achieves the fastest results, while `ralphi`’s runtime is comparable with WhatsHap at lower coverage and faster at higher coverage. On short reads, HapCUT2 is the fastest, with `ralphi` achieving the second best running time (note: we do not downsample short reads, which accounts for the increase in runtime at higher coverages). We also show `ralphi`’s total peak memory usage (the maximum resident set size) at each coverage. Since `ralphi`’s model is very small, `ralphi` is able to achieve a competitive runtime and a low memory footprint even when running with a single thread.

**Table 1:** Runtime (min) for each tool on chr1 of the NA12878 genome with ONT long reads. Memory usage is reported for `ralphi`.

| Tool | 5x | 10x | 15x | 30x |
| --- | --- | --- | --- | --- |
| HapCUT2 | 0.48 | 0.97 | 1.47 | 2.34 |
| LongPhase | 0.28 | 0.47 | 0.82 | 1.08 |
| RefHap | 86 | NA | NA | NA |
| WhatsHap | 1.00 | 3.00 | 7.44 | 10.3 |
| **ralphi** | 1.38 | 3.45 | 6.57 | 7.53 |
| Memory (GB) | 1.2 | 2.03 | 2.9 | 3.6 |

**Table 2:** Runtime (min) for each tool on chr1 of the NA12878 genome with Illumina short reads. Memory usage is reported for `ralphi`.

| Tool | 5x | 10x | 15x | 30x |
| --- | --- | --- | --- | --- |
| HapCUT2 | 0.23 | 0.45 | 0.65 | 1.29 |
| RefHap | 23.98 | 676.88 | NA | NA |
| WhatsHap | 1.84 | 5.35 | 7.71 | 12.17 |
| **ralphi** | 1.46 | 3.06 | 4.32 | 8.30 |
| Memory (GB) | 1.4 | 2.2 | 2.9 | 5.3 |

## 4 Conclusion

In this work we motivate the use of deep reinforcement learning for read-based haplotype assembly using the reduction of this task to the NP-hard MFC objective on fragment graphs. Our approach leverages (1) deep graph neural networks to learn effective graph state representations for complex and heterogeneous fragment graph topologies and (2) a reinforcement learning model to learn the optimal policy for partitioning the fragments into haplotypes while maximizing the MFC objective. Our results show that this approach can learn an effective strategy for haplotype assembly and outperforms the state of the art using inputs from different sequencing platforms. We plan to extend the framework to additional sequencing platforms and variant types (e.g., indels and SVs) in future releases.

## 5 Acknowledgments

This work was supported by the Broad Institute Schmidt Fellowship (V.P.), NIH R21HG013567 (V.P.), and NIH 1R35GM141861 (B.B.). We would like to thank the authors of WhatsHap for providing a robust and well-documented software framework, which allowed us to easily use and extend its read selection and realignment modules for pre-processing.

## Notes

### Competing Interest Statement

The authors have declared no competing interest.

